# Bacterial alteration of redox stressors impact environmental stability of Influenza A virus

**DOI:** 10.1101/2024.10.31.621345

**Authors:** Matthew Williams, Hannah M. Rowe

## Abstract

Influenza A virus (IAV) causes annual morbidity and mortality and remains a constant pandemic threat due to emergence of novel strains. Therefore, understanding the factors important in host-to-host transmission of IAV is a key control point for protecting individual and public health. Transmission is highly heterogeneous with viral factors and host inflammatory and immune factors being implicated. Also implicated is the upper respiratory microbiome. While typically thought to act indirectly on viral pathogenesis, in an immunomodulatory capacity to enhance or reduce susceptibility to viral infection, recent studies on the pathogenesis of IAV have identified direct interactions between the virus and upper respiratory pathobiont bacteria. We hypothesize that the bacterial cells and their metabolites co-shed into respiratory droplets with IAV particles, can alter the viability of the IAV particles in the environment, and therefore altering the capacity for host-to-host transmission. In this investigation we utilize a simplified model of fomite transmission in the absence of confounding host factors and demonstrate how oxidative stress from both the environment and the metabolic activity of *S. pneumoniae* contribute to the killing of IAV, while catalase or the metabolic activity of *S. aureus* can protect IAV from environmental or pneumococcally-produced reactive oxygen species.

## Introduction

Transmission of infectious disease is a complex process by which an infected host sheds a pathogen and then the pathogen must retain viability and infectivity until it encounters a new susceptible host, and then successfully establish infection in the new host. Understanding the control points in pathogen transmission is key for protecting individual and public health. One pathogen of great historical and current public health threat is Influenza A virus. While vaccination has decreased morbidity and mortality(1), current strategies are insufficient to block transmission(2, 3). Influenza A viruses are shed and acquired from the upper respiratory tract(4) in humans. The major mechanisms of transmission include both airborne particles that are directly inhaled(5-8) by the new host, and fomite mediated transmission where the particles or respiratory secretions are deposited onto objects, and then the new host through hand to mucosa contact will self inoculate(9, 10), or alternatively the fomites can serve as sources for re-aerosolization(11) and inhalation infection.

During airborne transit, and particularly during fomite-mediated transmission, IAV particles are subject to environmental stressors that can damage the envelope, the viral proteins, or the viral RNA, rendering the particles less infectious or non-infectious. Temperature and humidity have been implicated in environmental survival of IAV(12-14). Further, the contents of the respiratory secretions have been shown to impact the time of viral survival(15). An additional component of the contents of respiratory secretions are bacterial cells resident in the human upper respiratory microbial community, as well as the products of bacterial metabolism. Microbial mediated control of transmission of IAV has been suggested, including in IAV household transmission cohorts(16-18), and in animal models(19, 20). The suggested mechanisms mainly are via immunomodulation, or through production of enhanced symptoms leading to expulsion of virus particles in the case of the infected host and a more receptive tissue environment in the case of the contact(19).

Additionally, recently described direct interactions have been reported between IAV and several upper respiratory pathobiont bacteria(21, 22). This suggests an additional bacterially mediated mechanism of modulating viral transmission through alteration of environmental survival through direct interaction with stabilizing or destabilizing bacterial components, or through the bacterial metabolites secreted into the immediate micro-environment stabilizing or destabilizing the adherent viral particles. We have previously shown that bacterial partners with polysaccharide capsules were shown to be one mechanism by which IAV particles can survive desiccation mediated stress and maintain viability and infectivity(20). Individual to individual variations between the upper respiratory tract microbial communities in their structural and functional composition could therefore allow different combinations of bacterial-viral complexes to be generated during viral shedding, which would impact environmental survival of IAV and therefore transmission potential.

The impact of direct interactions between bacterial cells and components and viral particles being important for viral transmission is not unprecedented. Several enteric virus families have been shown to directly bind to bacterial cells(23-28) and to interact with bacterial cell wall carbohydrates(23, 26, 28). These interactions have been shown to improve viral infectivity by enhancing receptor engagement(26, 27). Further, these interactions have been shown to protect the stability of the virions in the environment(23, 26), allowing them to maintain infectivity between hosts, and therefore increase transmission potential.

Microbiome analyses often focus on a “who” model, characterizing the presence or absence of certain members of a microbial community and associating that with microbial co-occurrence patterns and health and disease states of the hosts. Expansion of community analysis to metagenomics, metatranscriptomics, and metabolomics, a “what are they doing” model, is characterizing what metabolic features of a community are important for microbial co-occurrence patterns and for health and disease states of the hosts. Here, we expand the “what are they doing” framework to describe the bacterial production of metabolites that could be both beneficial or harmful to co-infecting viruses. In this work, we focus on one metabolic feature of microbes found in the upper respiratory microbial community, that of production and detoxification of reactive oxygen species, during normal bacterial metabolism and bacterial stress responses. This then leads to a model of microbiome-mediated IAV transmission, where following shedding of virions from the infected host, these virions are in the context of the microbial cells when they are introduced into the environment. During environmental transit, ongoing metabolism by the bacteria would be producing products that could protect or could damage the viral particles, altering the ability for those particles to be infectious upon encountering a new host. Once such bacterial metabolic reaction could be producing reactive oxygen species and destabilizing the viral particles-reducing transmission potential, or ongoing metabolism by the bacteria could be detoxifying reactive oxygen species produced spontaneously in the environment, enhancing virion stability-increasing transmission potential.

In this work we utilize a simplified model of IAV fomite mediated survival, with no host-derived infection signals. We demonstrate that metabolites produced by respiratory pathobionts including *S. pneumoniae* can reduce survival of IAV in the environment, while metabolites produced by other respiratory pathobionts, including *S. aureus*, can protect IAV from environmentally-mediated viability loss. Further, the factors produced by *S. aureus* can protect the viral particles from the factors produced by *S. pneumoniae* suggesting polymicrobial impacts on IAV environmental stability and therefore transmissibility.

## Results

In previous studies(20, 22, 29) we produced bacterial-viral complexes through pre-mixing bacterial cells and viral particles and then subjected these complexes to rapid desiccation using vacuum and centrifugation in a speedvac. This work showed that some, but not all, bacterial species found in the human upper respiratory microbial community could protect IAV particles from desiccation mediated viability loss(20), and that this seemed to be at least partially explained by bacterial expression of polysaccharide capsule. Here we wished to further investigate the impacts of bacterial cells and components on desiccation mediated viability loss of IAV. However, we wished to perform a more gentle and biologically relevant desiccation. Therefore we mixed washed bacterial cultures resuspended in phosphate buffered saline (PBS) with viral particles and allowed them to desiccate in normal room conditions for 24 hours in the dark. Using this more gentle desiccation protocol, we were able to see environmental survival of IAV. We were surprised to see that *Streptococcus pneumoniae*, which improved viral survival under the rapid desiccation protocol(20), in fact reduced the survival of IAV (Figure 1A). Also, *Staphylococcus aureus*, which provided minimal protection of IAV from rapid desiccation in our previous model(20), was able to significantly increase viral survival compared to virus alone (Figure 1A). The impact of typical Gram negative pathobionts, *Moraxella catarrhalis* and non-typable *Hemophilus influenzae* on viral survival was less profound. *M. catarrhalis* provided significant protection, concurring with our prior findings in the harsher desiccation conditions, while *H. influenzae* did not protect, nor did it kill IAV particles, again concurring with prior findings(20).

**Figure 1:**
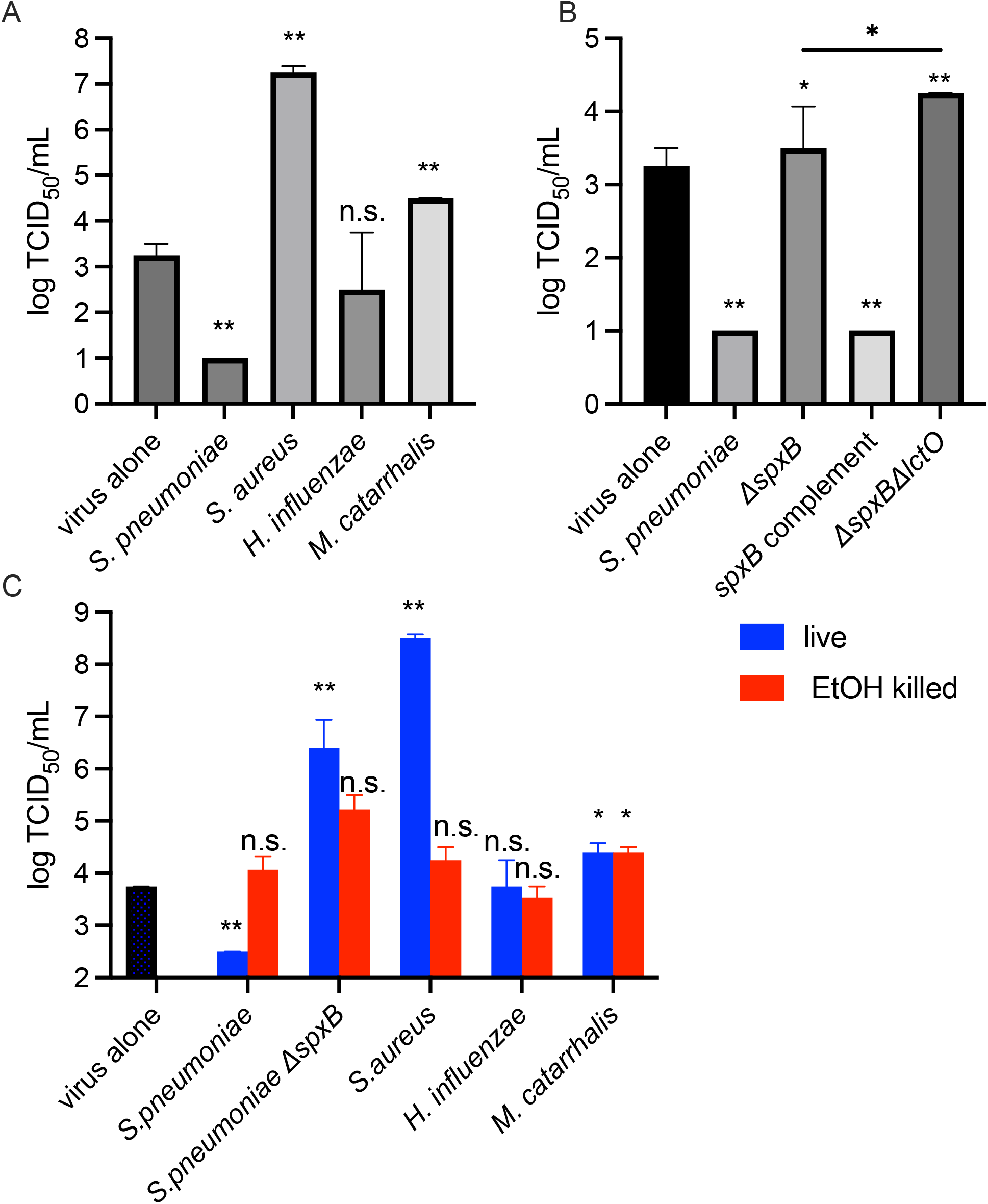
Bacterial H_2_O_2_ production and detoxification mediate viral environmental survival and requires live bacteria. **A**. 5 log TCID_50_ IAV strain PR8 was desiccated at room temperature for 24 hours alone or in the presence of 6 log indicated bacterial strain in PBS. **B**.5 log TCID_50_ IAV strain PR8 was desiccated at room temperature for 24 hours alone or in the presence of 6 log live or ethanol killed bacteria of indicated species. **C**. 5 log TCID_50_ IAV strain PR8 was desiccated at room temperature for 24 hours alone or in the presence of 6 log wild type *S. pneumoniae*, or isogenic mutants of the hydrogen peroxide producing enzymes of *S. pneumoniae: spxB, lctO*, or double mutant. Viable virus after desiccation was determined by TCID_50_ on Manin Darby Canine Kidney cells. Data are plotted as median with error bars indicating interquartile range. \**p<0.05, **p<0.01*, n.s. not significant by Mann Whitney.

To determine whether the modulation of viral viability is the result of bacterial metabolic activity, or intrinsic to the bacterial cells, we next desiccated the virus in the presence of ethanol-fixed *S. pneumoniae, S. aureus, H. influenzae* or *M. catarrhalis*. We had previously shown that ethanol-fixed cells maintained the ability to interact with IAV particles to form the bacterial-viral complex(22). When desiccated in the presence of ethanol fixed *S. pneumoniae* or *S. aureus*, IAV retained equivalent infectivity to virus alone (Figure 1B), while live bacteria significantly altered the survival of the virus under desiccation stress compared to killed bacteria. This suggests that it is not a component intrinsic to these bacterial cells that alters IAV desiccation tolerance, but a product of active metabolism by the bacteria. *H. influenzae* cells, live or ethanol killed, did not promote or diminish environmental survival of IAV, while both live and ethanol killed *M. catarrhalis* both promoted viral survival, suggesting that an intrinsic component of the *M. catarrhalis* cell is capable of promoting viral environmental survival.

Previously(29) we had shown that *spxB* mutant *S. pneumoniae* had a defect in both host-to-host bacterial transmission and in environmental survival of *S. pneumoniae*. SpxB, pyruvate oxidase, is a key player in pneumococcal central metabolism(30), and has roles in modulating capsule levels(31, 32). Most critically for mediation of environmental survival is through the production of the reactive oxygen species hydrogen peroxide (H_2_O_2_). As pneumococci do not possess a catalase, they cannot detoxify this metabolite and are subject to oxidative damage and loss of viability. In our previous work(20) using speedvac mediated desiccation, *spxB* mutant pneumococci were shown to be better able to protect IAV than wild type pneumococci from desiccation-mediated viability loss, suggesting H_2_O_2_ production by *S. pneumoniae* is destabilizing the virions. Next we tested the *spxB* strain in our gentle desiccation assay for its ability to impact IAV environmental persistence. Here we show no desiccation-mediated viability loss when IAV is desiccated in the presence of *spxB* mutant *S. pneumoniae* (Figure 1C) which is reversed in the *spxB* complemented strain, supporting a role for H_2_O_2_ production by pneumococci in damaging IAV particles and reducing environmental survival. Pneumococcal lactate oxidase, LctO, also produces hydrogen peroxide as a byproduct of metabolism(33). To test the impact of further reducing the ability of the pneumococcus to produce hydrogen peroxide, we next made a strain with both *spxB* and *lctO* deleted and examined the ability for this strain to protect IAV in our model of gentle desiccation. The *spxB lctO* double deletion strain demonstrated significantly higher protection to the IAV particles compared to the *spxB* single deletion (Figure 1C).

As we previously showed bacterial capsule, in a rapid model of desiccation, to be important for IAV desiccation tolerance(20), and *spxB* deficient pneumococci exhibit altered capsule production(31), we next asked if ethanol-fixed *spxB* mutant pneumococci were capable of stabilizing IAV against desiccation stress in our model of gentle desiccation as well as live *spxB* mutant bacteria could (Figure 1B). There was no significant difference in viral stability when desiccated in the presence of live versus killed *spxB* mutant bacteria, suggesting that the lack of metabolic production of H_2_O_2_ is sufficient for the altered viral desiccation tolerance.

While *S. pneumoniae* does not produce a catalase to detoxify H_2_O_2_, many bacterial species do, including *S. aureus*. We next asked if *S. aureus* could protect IAV from desiccation mediated viability loss in the presence of wild type, H_2_O_2_ producing, *S. pneumoniae*. When IAV was desiccated in the presence of live wild type *S. pneumoniae* and live *S. aureus*, IAV particles were protected from viability loss (Figure 2A), suggesting a role for the staphylococcal catalase in detoxification of pneumococcal hydrogen peroxide. To determine whether metabolic activity of both bacterial species was required for alteration of viral environmental stability, we desiccated the virus in the presence of a mix of live and ethanol-fixed *S. aureus* and *S. pneumoniae* cells. When live *S. aureus* was present, regardless of if the pneumococci were live or killed, the virus was significantly protected from viability loss compared to virus alone (Figure 2A), suggesting that staphylococcal metabolism is required for protecting the virus from environmental mediated viability loss. When live pneumococci were present, they required live *S. aureus* to be present for the virus to retain viability, suggest that a staphylococcal metabolite was necessary for protecting the virus from pneumococcal metabolites, and not merely the presence of staphylococcal cells to protect the virus from pneumococcal metabolites. Similarly, when the virus was desiccated in the presence of killed bacteria from both species, there was no significant difference compared to virus alone, supporting the role of bacterial metabolites in protecting the virus from environmental mediated viability loss.

**Figure 2:**
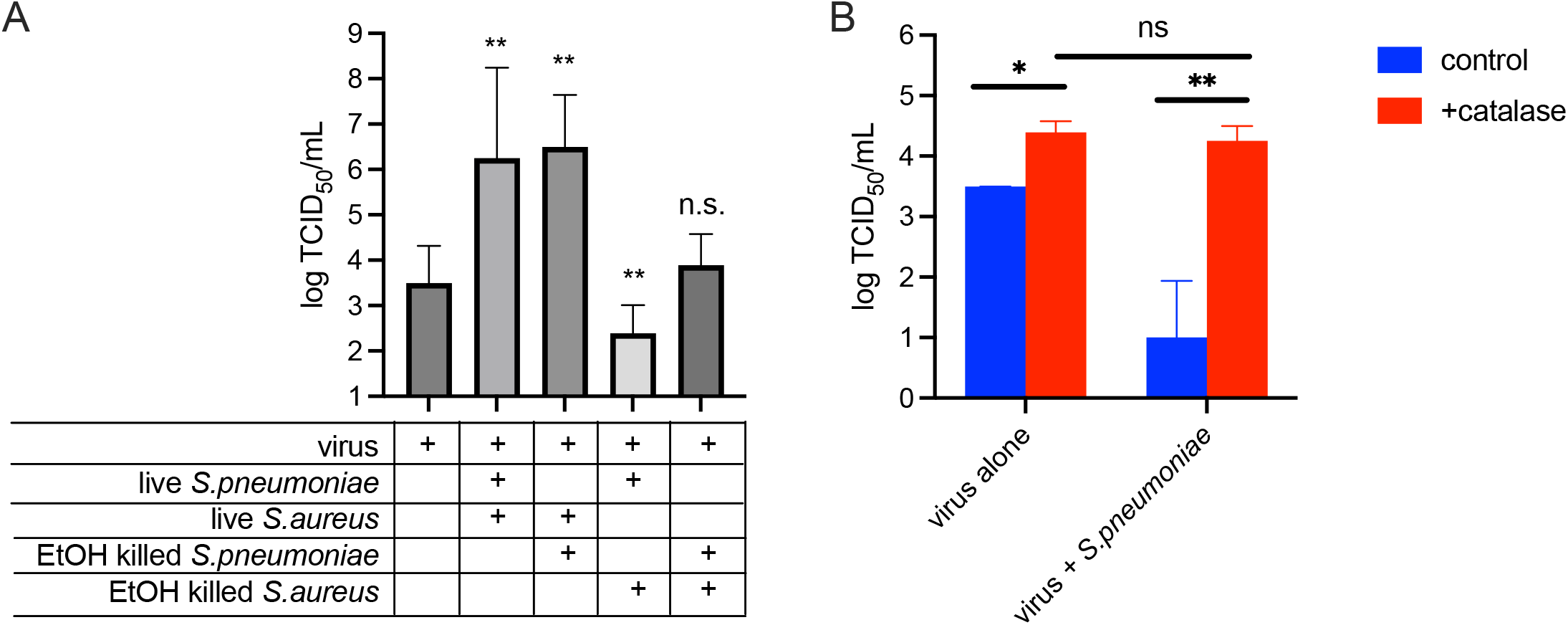
Extracelluar catalase activity can protect IAV from pneumococcally generated and environmental ROS. **A**. 5 log TCID_50_ IAV strain PR8 was desiccated at room temperature for 24 hours alone or in the presence of 6 log indicated bacterial strain in PBS. **B**. 5 log TCID_50_ IAV strain PR8 was desiccated at room temperature for 24 hours alone or in the presence of 6 log wild type *S. pneumoniae*, in PBS (control) or 0.01% catalase. Viable virus after desiccation was determined by TCID_50_ on Manin Darby Canine Kidney cells. Data are plotted as median with error bars indicating interquartile range. \**p<0.05* by Mann Whitney.

To further confirm the roles of catalase in protecting IAV particles from environmental and bacterially-produced reactive oxygen species, we repeated the desiccation protocol in the presence of 0.01% catalase. Catalase alone was able to provide significant protection against viability loss igure 2B). Further, catalase was able to reverse the impact of *S. pneumoniae*, and allow the virus to survive in the presence of live pneumococci, supporting a role that H_2_O_2_ produced by pneumococci is destabilizing IAV.

## Discussion

A key knowledge gap in understanding and controlling influenza transmission is understanding the factors influencing environmental stability. While temperature, humidity and IAV strain factors have been identified, these cannot fully explain the heterogeneity of transmission. Another possible factor could be the human-to-human variability in the upper respiratory microbial community. Our results support a model whereby during shedding from an infected human upper respiratory tract, IAV particles are deposited on a surface in the context of metabolically active bacteria. The results shown here demonstrate that bacterial metabolism can modulate the reactive oxygen species environment and therefore alter the environmental survival of the virus and ability to transmit host-to-host through airborne, re-aerosolization or fomite-mediated transmission. We also anticipate that bacterial metabolites and bacterial cell components could also be operative in altering exposure of viral particles to environmental stresses other than reactive oxygen species, though with less dramatic impact. The individual-to-individual variations present in the density and composition of the microbial community could in part explain the heterogeneity of transmissibility.

The work reported here is in contrast to other reports(34) that show that *S. pneumoniae* is capable of protecting IAV during desiccation in the presence of host mucus, however, catalase is present in mucus, and therefore these results do not disagree. We wanted to isolate the bacterial factors to examine their impact outside of the confounding variable of host factors. Our work also contrasts with another recent study(35) showed that pneumococcus can protect IAV in the environment using a saline solution without confounding host factors, however, this study looked at earlier time points and smaller volumes, modeling a different transmission mode (quicker drying respiratory droplets instead of longer drying fomites), and therefore different mechanisms are likely at play, with pneumococcus being protective in the biophysics of the droplet preventing salt supersaturation killing of IAV(36), but later time points in larger volumes, peroxide production damaging the virions and reducing viral environmental survival.

This study is also limited by only using a single viral strain, as the environmental stability of IAV strains varies(14) we might anticipate other viral strains to exhibit more or less sensitivity to bacterial reactive oxygen species and other bacterial metabolites. Our prior work with harsh desiccation did show differences in the magnitude of protection of H3N2 IAV strains compared to H1N1 strains(20), supporting different interactions between viral strains and bacterial cells and metabolites.

## Materials and Methods

### Bacterial cultures

*S. pneumoniae* strain BHN97 and derivatives were grown from freezer stock on solid media: Tryptic Soy Agar (BD) supplemented with 3% defibrinated sheep blood (Lampire) in a humidified 5% CO2 incubator at 37 degrees Celsius. Growth from plates was transferred to THY Broth: Todd Hewitt (BD) + 0.2% yeast extract (VWR), and was grown static in glass tubes in a humidified 5% CO2 incubator at 37 degrees Celsius until mid-log, OD_620_ approximately 0.4. BHN97 *spxB* and *spxB-*complemented strain were generated previously(29). BHN97 *spxBlctO* mutant was made for this study (see below.) *S. aureus* strain MW2 (gift from Alex Horswill) was grown from freezer stock in liquid Brain Heart Infusion (BD) (BHI) media overnight at 37 degrees Celsius with aeration. Overnight cultures were diluted 1:200 in fresh BHI and grown at 37 degrees Celsius with aeration until mid-log, OD_620_ approximately 0.4. Non-typeable *H. influenzae* strain 86–028NP was grown on chocolate agar (Remel) and then directly inoculated into brain heart infusion broth supplemented with 0.2% yeast extract, 10 μg/mL hemin (Sigma) and 10 μg/mL NAD (Sigma) and grown with aeration to the mid-log phase, OD_600_ approximately 0.18(37). *M. catarrhalis(38)* was grown on non-supplemented TSA plates, directly inoculated in BHI broth supplemented with 0.2% yeast extract and grown with aeration to the mid-log phase, OD_620_ approximately 0.4.

### Ethanol killed bacteria

Ethanol killed bacteria were made as previously described(22). Briefly, 10^8^ CFU mid log culture was resuspended in ice cold 70% ethanol and allowed to incubate on ice 5 minutes. Bacteria were pelleted, ethanol was aspirated, and killed bacteria were resuspended in 1 mL PBS. Killing was verified by plating 100uL of killed material on TSA/3% sheep blood and overnight incubation and detection of zero colonies.

### Making *spxBlctoO* double deletion

Allelic exchange with a spectinomycin resistance cassette was performed as previously described(22). About 1kb upstream of *lctO* was amplified from BHN97 genomic DNA using primers (BHN97 LctO up F-GATGGCCCAGTCAACTACC and BHN97 LctO up R spect-TGTATTCACGAACGAAAATCGATAAAATGCCCTCCTTGATTAAGTAAG) and about 1kb downstream of *lctO* was amplified with primers (BHN97 LctO down F spect-GAAAACAATAAACCCTTGCATATGTAAAACAGATTGCCTCCACTGAATG and BHN97 LctO down R-GCGACTGCTTGATTCCAGC) using PrimeStar High Fidelity Polymerase (Takara). PCR products were gel extracted (Qiagen MinElute) and along with the spectinomycin resistance cassette were combined using splicing by overlap extension PCR using PrimeStar High Fidelity Polymerase (Takara). Purified PCR product was transformed into *spxB* deletion strain of BHN97, in THY media supplemented with 0.002% BSA, 0.2% glucose and 0.0002% CaCl_2_ (39) grown to early log (OD_620_ approximately 0.2), diluted 1:100 in supplemented THY, incubated 14 minutes with 2ul/mL each 1mg/mL CSP-1 and CSP-2(40).

### Cell culture

Manin Darby Canine Kidney (MDCK) (ATCC CCL-34) were grown in MEM (Gibco) supplemented with 10% Fetal bovine serum (Cytiva), 1x Glutamax (Gibco), and 1x Sodium Pyruvate (Gibco), in a humidified 5% CO2 atmosphere at 37 degrees Celsius.

### Viral cultures

Influenza strain A/Puerto Rico/08/1934 (H1N1) was grown in greater than 90% confluent MDCK cells in Infection Media: MEM (Gibco) supplemented with 1X Gluatmax (Gibco), 0.075% Bovine Serum Albumin Fraction V (Gibco), and 1ug/mL TPCK-Trypsin (Worthington), for 96 hours in a humidified incubator with 5% CO2 atmosphere at 37 degrees Celsius. Culture supernatant was clarified by centrifugation at 500 x*g* for 5 minutes and was aliquoted into 0.5mL volumes and stored at -70 degrees Celsius.

## Desiccation

Bacterial strains were grown to mid-log phase and normalized to about 10^8^ CFU/mL in PBS. 6.2 log TCID_50_ A/Puerto Rico/08/1934 in a volume of 50uL was added to 24 well plates. 100 uL bacterial culture, or PBS for virus only controls, was added to each well and swirled to mix. When two bacterial strains were mixed, 50uL of each bacterial strain was used, for a total of 10^7^ CFU and 100 uL volume of bacterial cells. Plates were left open in the dark at room temperature overnight. Dessicated material was resuspended in 10X suggested working concentration of PenStrep (1000 U/mL Penicillin, 1000 U/mL Streptomycin) solution (Gibco) diluted in sterile de-ionized water and stored at -70 degrees Celsius.

Catalase from Bovine Liver (Sigma) was reconstituted and diluted in PBS so that the final concentration, after addition of bacterial cells and viral culture was 0.01%. 6.2 log TCID50 A/Puerto Rico/08/1934 in a volume of 50uL was added to 24 well plates. 50uL bacterial culture, or PBS for virus only controls, and 50uL of PBS or catalase, was added to each well and swirled to mix, and desiccated and resuspended as above.

### Determination of viable virus following desiccation

Rehydrated material was thawed at 4 degrees Celsius. Viral titer was determined by 50 percent tissue culture infectious dose assay as described previously(22). Briefly, MDCK cells were seeded at 3×10^4^ cells per well in 96 well plates. Serial 10 fold dilutions of rehydrated samples were made in Infection media. Cells were infected in triplicate with each dilution. 72 hours post infection, supernatant from infected cells was mixed 1:1 with 0.5% turkey red blood cells in PBS (Lampire) in V bottom 96 well plates for hemagglutination assay. Plates were incubated at room temperature for 30 minutes. Hemagglutination was read for each well and 50% tissue culture infectious dose was calculated using the method of Reed and Munch(41).

Statistical analyses were performed using GraphPad Prism Version 10.3.1. Data was tested for normality with the Kolmogorov-Smirnov test and found to have non-normal distribution. Therefore the Mann-Whitney test was performed for pairwise comparisons.

